# Medium-length stamen mediates delayed selfing in a two-step self-pollination system in *Commelina communis*

**DOI:** 10.64898/2026.04.02.716232

**Authors:** Kaho Ikeda, Yoshiaki Kameyama

**Affiliations:** Faculty of Regional Environment Science, Tokyo University of Agriculture, Tokyo 156-8502, Japan

**Keywords:** *Commelina communis*, heteranthery, pollen movement, quantum dots, two-step self-pollination

## Abstract

We examined the functional roles of trimorphic stamens in self-pollination in *Commelina communis* using quantum dots to label pollen grains. In the absence of pollinators (i.e., under greenhouse conditions), bud pollination and delayed autonomous selfing contributed equally to self-pollination. Delayed autonomous selfing was achieved primarily by the anther of the medium-length stamen (M-anther), whereas the long anthers (L-anthers), positioned adjacent to the stigma during anthesis, and the short anthers (S-anthers), which produce a small number of infertile pollen grains, had no contribution. In field experiments, a low pollinator visit frequency restricted outcross pollination, whereas L-anthers increased their contribution to self-pollination to levels comparable to those of bud pollination and M-anther–mediated selfing. Our results reframe heteranthery in *C. communis* as part of a temporally structured two-step selfing system and provide a basis for reassessing its functional significance under variable pollination environments.

## 1. INTRODUCTION

Angiosperms have evolved diverse floral traits and mating systems in response to variations in pollination environments. Floral morphology often reflects functional adaptations associated with mating strategy. For example, heterostyly involves the reciprocal placement of anthers and stigmas among floral morphs and promotes disassortative mating while reducing selfing and sexual interference (Barrett, 2002; Darwin, 1877; Lloyd & Webb, 1992). Conversely, predominantly selfing species commonly exhibit small flower size, reduced herkogamy, and lower pollen-to-ovule ratios, traits associated with reduced pollinator dependence (Barrett, 2002; Harder & Johnson, 2023; Heywood et al., 2022; Tsuchimatsu & Fujii, 2022).

Many hermaphroditic species exhibit mixed mating (i.e., a combination of selfing and outcrossing), and the rate of selfing can vary widely among closely related species and even among populations within species (Goodwillie, 2005; Kalisz, 2004; Kameyama & Kudo, 2015; Leibman et al., 2018). The timing of selfing can be categorized as prior, competing, or delayed selfing based on when self-pollen is deposited relative to outcross pollen (Goodwillie, 2005; Lloyd, 1979; Lloyd & Shoen, 1992). Two-step self-pollination is a reproductive strategy in which prior selfing is followed by delayed selfing while retaining the opportunity for outcrossing, thereby potentially enhancing reproductive assurance under adverse conditions (Domingos-Melo et al., 2018; Liu et al., 2025).

In species with morphologically differentiated stamens, temporal structuring of selfing might be associated with functional differentiation among stamen types. Heteranthery, describing the presence of morphologically distinct stamens within a flower, has often been discussed in the context of a division-of-labor hypothesis, in which different anthers serve feeding and pollinating functions (Mesquita-Nato et al., 2017; Tang & Huang, 2007; Vallejo-Marín et al., 2010). However, most studies have focused on its implications for outcrossing, and its potential role in structuring different modes or timing of selfing remains largely unexplored. Quantitative evaluation of pollen deposition by distinct stamen types is therefore essential for understanding the potential contributions of heteranthery to mating dynamics.

*Commelina communis* is an annual, andromonoecious herb native to temperate northeastern Asia (Figure 1). Plants bear many inflorescences, each producing a few short-lived flowers per day that open at sunrise and close at noon. Flowers have three types of stamens: two long brown stamens (L-anthers), one medium-length yellow stamen (M-anther), and three short yellow stamens (S-anthers) that produce small numbers of infertile pollen grains and function as staminodes (Morita & Nigorikawa, 1999). The stigma is positioned at approximately the same height as the L-anthers. At the end of the anthesis, the pistil and L-anthers coil, presumably facilitating autonomous self-pollination (Morita & Nigorikawa, 1999). Syrphid flies (e.g., *Episyrphus balteatus*) are the primary pollinators, although bumblebees, honeybees, and small halictid bees also visit the flowers (Ushimaru & Hyodo, 2005; Ushimaru et al., 2007, 2014). Pollen serves as the sole reward for pollinators.

**Figure 1.**
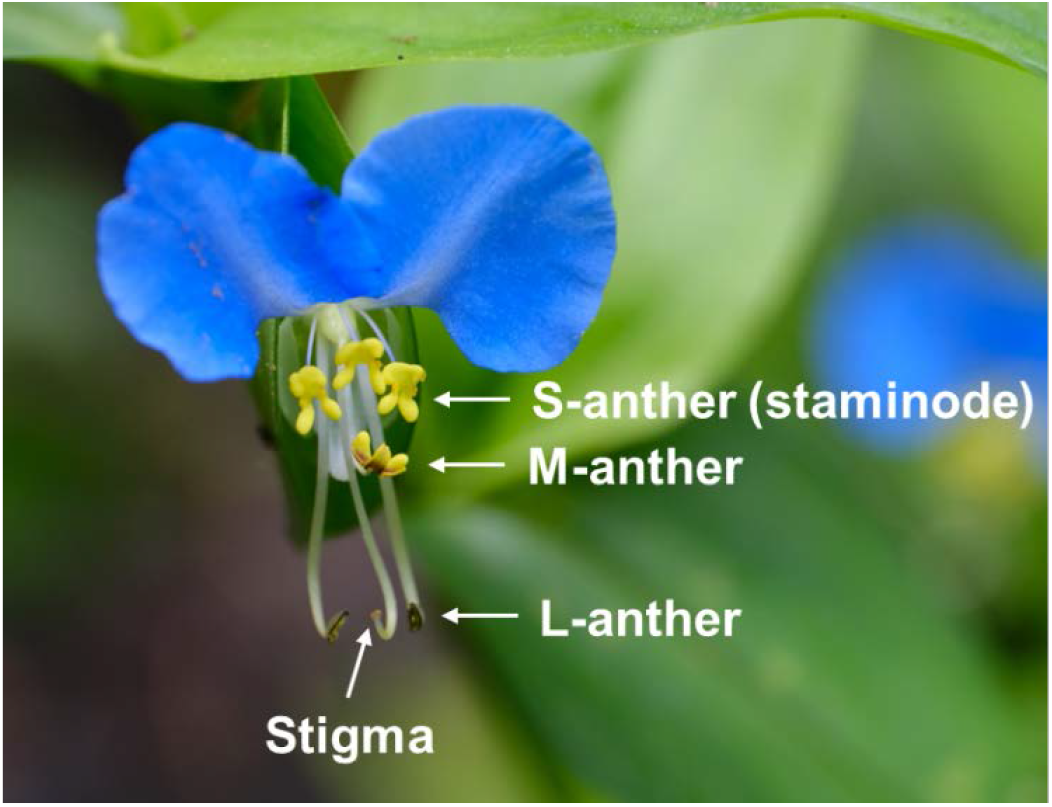
Perfect flower of *Commelina communis*. Perfect flowers bear two long brown stamens (L-anthers), one medium-length yellow stamen (M-anther), and three short yellow stamens (S-anthers), which produce small numbers of infertile pollen grains and function as staminodes. The stigma is positioned at approximately the same height as the L-anthers.

In *C. communis* L. (Commelinaceae), which possesses trimorphic stamens and exhibits both bud pollination and delayed autonomous selfing (Katsuhara & Ushimaru, 2019; Morita & Nigorikawa, 1999), the relative contribution of each stamen type to selfing remains unclear. In this study, we partitioned the functional roles of L-, M-, and S-anthers to assess the integration of heteranthery into a two-step self-pollination system.

## 2. MATERIALS AND METHODS

### 2.1 Study sites and sample collection

In May 2025, 30 *C. communis* individuals were collected from three sites (10 individuals per site) on the campus of Tokyo University of Agriculture (Tokyo, Japan; 35.64126° N, 139.63215° E). The sites were separated by several tens to a few hundred meters. Each plant were transplanted into a plastic slit pot (136 mm in height, 115 mm in diameter). A 2–3-cm layer of pumice was placed at the bottom of each pot for drainage, and the pots were filled with red soil amended with a slow-release fertilizer (MagAmp K, Hyponex Co., Ltd., Osaka, Japan). The plants were grown in a sunny field until the experiments were conducted.

### 2.2 Greenhouse experiments

To quantify pollen production, we arbitrarily selected five perfect flowers (one per individual) immediately before anthesis and collected L-, M-, and S-anthers. Anthers of each type from each flower were stored separately in 500 μL of 70% ethanol. The total number of pollen grains per anther type was estimated by counting pollen grains in three 5.0-μL subsamples per sample under a microscope (×100).

To quantify pollen movement in the absence of pollinators (i.e., under greenhouse conditions), we arbitrarily selected 30 perfect flowers (2–3/individual). Immediately after anthesis, pollen grains of the L- and M-anthers were labeled with differently colored quantum dots (red and green, respectively; Strem Chemicals, Newburyport, MA, USA; see Minnaar & Anderson 2019 for detailed protocols). Quantum dots were dissolved in hexane (5 mg/mL) and applied to anthers in 2-μL volumes per anther. Stigmas were collected after flower closure and mounted on microscope slides in glycerin jelly (5 g of gelatin, 40 mL of glycerin, and 30 mL of distilled water) containing 1 mL of iodine solution as a preservative. Pollen grains on the stigma surface were then counted under a microscope (×100) with 365-nm UV light. Unlabeled pollen represented bud pollination, whereas labeled red and green pollen represented delayed selfing from the L- and M-anthers, respectively. The experiments were conducted in August 2025.

### 2.3 Field experiments

To quantify pollen movement in the field, we established two artificial plots and one natural plot (approximately 1.5–2.5 m^2^), each separated by several tens of meters. In the artificial plots, five potted individuals were arranged, each bearing one or two perfect flowers (mostly two). In the natural plot, extra flowers were removed to standardize flower numbers. Overall, each plot contained approximately 9–10 perfect flowers (mean = 9.8). Immediately after anthesis, pollen grains of the L- and M-anthers of a single target flower were labeled with quantum dots. The stigmas of all flowers were collected after flower closure, and pollen grains on the stigma surface were observed under a microscope. Unlabeled pollen represented a mixture of bud pollination and outcrossing with nontarget flowers. On the stigmas of target flowers, red and green pollen represented delayed selfing from the L- and M-anthers, respectively. On the stigmas of nontarget flowers, red and green pollen represented outcrossing from the L- and M-anthers of the target flower, respectively.

Concurrently, pollinator visitation was observed within each plot for 15 min at five hourly intervals (6:00–11:00). Pollinator visit frequency (mean number of visits per flower per day) was calculated as follows: visit frequency = (total number of visits recorded across the five 15-min observation periods × 4)/number of flowers in the plot, where the factor of 4 accounts for the 15-min sampling within each hour of the flowering period. The experiments were conducted from August 1 to October 23, 2025 (35 trials over 15 days).

### 2.4 Statistical analysis

In the field experiments, pollen deposition was recorded on both target and nontarget flowers. However, all statistical analyses focused on target flowers, as labeled pollen was rarely detected on nontarget stigmas (see Supplementary Figure S1). Similarly, unlabeled pollen of target flowers could theoretically originate from either bud pollination or outcrossing, but in practice, almost all unlabeled pollen originated from bud pollination. Therefore, unlabeled pollen on target flowers was treated as bud pollen in both greenhouse and field experiments.

The number of pollen grains on the stigma surface was analyzed using a generalized linear mixed model with a negative binomial error distribution (nbinom2). Pollen category (unlabeled, red-labeled, and green-labeled) and experiment (greenhouse vs. field), as well as their interaction, were included as fixed effects. To account for the hierarchical structure of the field data, the plot and date nested within the plot were included as random effects for environmental hierarchy, whereas flower identity was included as a separate random effect to account for variation among flowers. Estimated marginal means were calculated for each combination of pollen category and experiment on the response scale, and pairwise comparisons were performed using Tukey’s adjustment for multiple testing. All analyses were conducted with *R* 4.5.2 (R Core Team, 2025) using the packages glmmTMB and emmeans. Unless otherwise indicated, data were presented as the mean ± SD.

## 3. RESULTS

### 3.1 Pollen production

Pollen production varied among stamen types. L-anthers produced the most pollen (4,766.7 ± 395.1), followed by M-anther (1,520.0 ± 65.0) and S-anthers (180.0 ± 29.8).

### 3.2 Pollen movement

In the greenhouse, stigmas received mostly unlabeled pollen (4.7 ± 4.7) and green pollen (3.3 ± 5.9), whereas red pollen was rare (0.4 ± 0.4; Figure 2). In the absence of pollinators, delayed autonomous selfing (green pollen) was achieved primarily by M-anther, and the levels of bud pollination (unlabeled pollen) and delayed selfing were comparable. L-anthers (red pollen) contributed almost no pollen.

**Figure 2.**
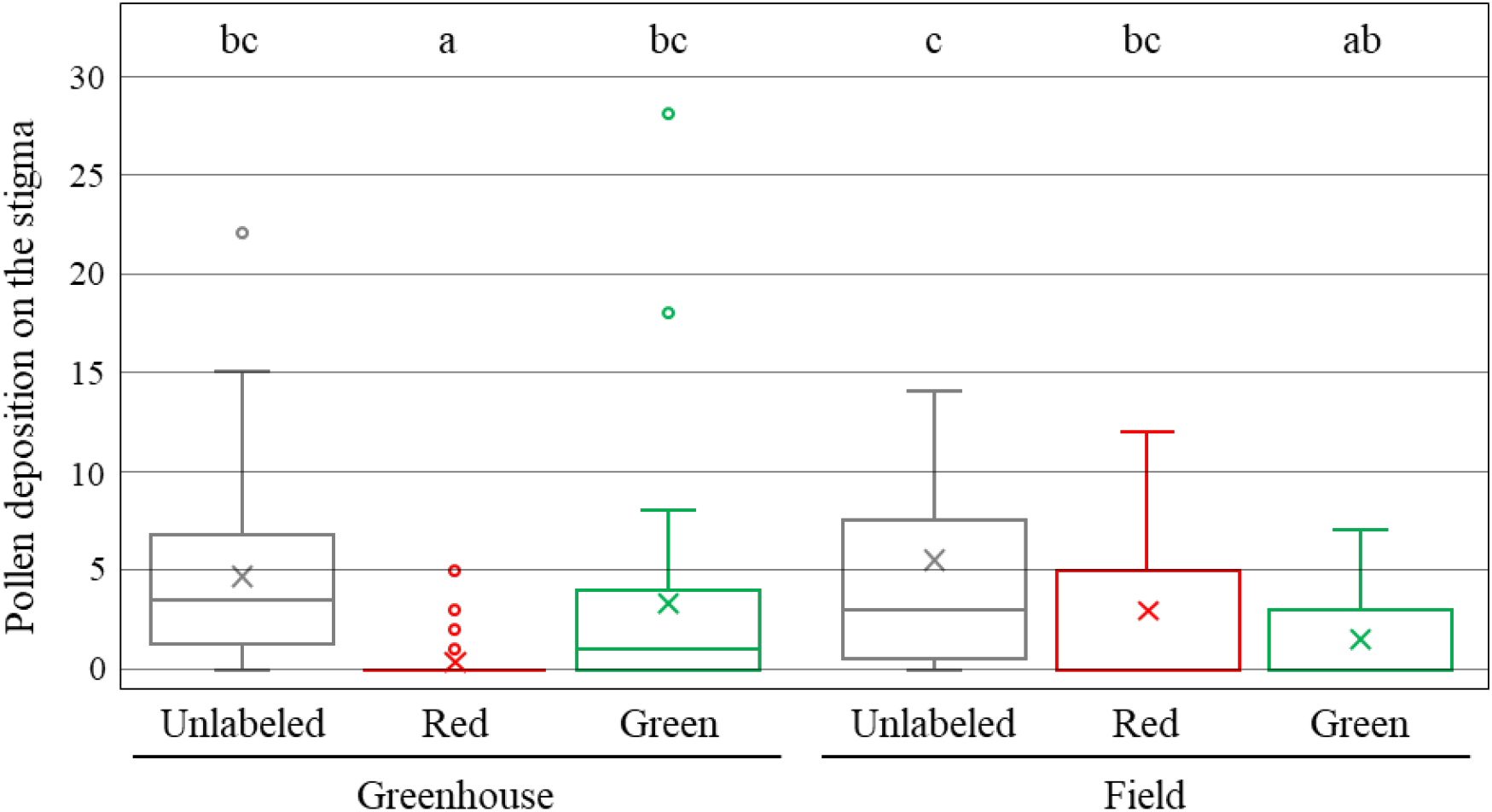
Pollen deposition on the stigmas of target flowers in greenhouse and field experiments. Unlabeled pollen represents bud pollination, whereas red and green labeling reflects long stamen and medium-length stamen origins, respectively. Only target flowers are presented. Boxplots show the median (horizontal line), the interquartile range (box), whiskers extending to 1.5 × interquartile range, and outliers (points). Crosses indicate means. Different letters indicate significant differences among groups based on Tukey-adjusted comparisons of estimated marginal means (*P* < 0.05).

In the field, target flowers received 5.5 ± 6.5 unlabeled, 3.0 ± 4.4 red, and 1.5 ± 2.6 green pollen grains (Figure 2), whereas nontarget flowers received almost exclusively unlabeled pollen (11.5 ± 9.1), with minimal red or green pollen detected (0.0 ± 0.3 and 0.0 ± 0.0, respectively), indicating that outcrossing from labeled target flowers was rare (Figure S1). In the presence of pollinators, the contribution of L-anthers to self-pollination increased significantly, reaching levels comparable to those of bud pollination and M-anther–mediated selfing.

### 3.3 Pollinator visitation

Among the 49 observed pollinators, most were small halictid bees (*Halictidae* sp.; 37 individuals [75.5%]). The remaining species included small and large syrphid flies, namely *Eumerus japonicus* (seven individuals [14.3%]) and *E. balteatus* (five individuals [10.2%]). The number of flowers visited per individual was 1.7, 1.6, and 3.0 for *Lasioglossum* sp., *E. japonicus*, and *E. balteatus*, respectively (overall mean = 1.8). The pollinator visit frequency per flower per day was 1.01 ± 1.22.

## 4. DISCUSSION

Heteranthery has long been interpreted in terms of the division of labor, with different anther morphs serving distinct roles in pollinator attraction and pollen export (Mesquita-Nato et al., 2017; Tang & Huang, 2007; Vallejo-Marín et al., 2010). Most studies have emphasized the role of this differentiation in promoting outcrossing. Alternative functional interpretations have been proposed (Dellinger et al., 2021; Kay et al., 2020; Telles et al., 2020), but they also have largely focused on cross-pollen transfer. Notably, Hrycan & Davis (2005) demonstrated in *C. dianthifolia* that M-anther can function as pollinator rewards and contribute to delayed selfing. Unlike *C. communis*, M-anther in that species produces more pollen than the L-anthers, and bud pollination has not been observed.

Two-step self-pollination allows plants to combine prior and delayed selfing while still retaining opportunities for outcrossing. Domingos-Melo et al. (2018) quantified their relative contributions in *Sida cordifolia*, revealing that they collectively account for most fruit production, whereas each alone is responsible for approximately half of that success. More recently, Liu et al. (2025) revealed the temporal and mechanistic basis of this strategy in *Arabidopsis*, demonstrating how spatially and temporally separated selfing events enhance fertilization efficiency under pollen-limited or stressful conditions. These studies provide a framework for understanding the ability of temporal separation to improve reproductive success and set the stage for clarifying the mechanism by which *C. communis* coordinates selfing through timing and anther specialization.

Our study of *C. communis* uncovered an underappreciated aspect of two-step self-pollination in a heterantherous floral system. Whereas previous research emphasized the temporal separation of anther function, we demonstrated that functional differentiation among anther morphs plays a critical role in regulating delayed selfing. M-anther contributes both as pollen rewards to promote legitimate pollination (Ushimaru et al., 2007) and as mediators of delayed selfing, with its contribution modulated according to pollinator availability. This flexible, context-dependent strategy preserves opportunities for outcrossing while ensuring reproductive assurance under pollinator limitation. By integrating temporal and structural mechanisms, floral architectures traditionally associated with promoting outcrossing can also facilitate adaptive selfing strategies, highlighting the evolutionary potential of anther specialization to balance multiple reproductive pathways within a single flower.

Such functional flexibility might also have ecological and evolutionary implications. Urban populations of *C. communis* exhibit decreased stigma height and reduced herkogamy, alongside a high fruit set, presumably reflecting reproductive assurance via selfing (Ushimaru et al., 2014). In addition, local adaptation to urban environments has led to divergence in key life-history traits, including flowering onset and plant size (Taichi et al., 2025). Investigating the interaction of anther differentiation with two-step selfing under these conditions could provide insights into the mechanisms underpinning reproductive success. Future studies could explore whether shifts in M-anther function, floral morphology, or the timing of selfing events contribute to these adaptive responses.

## ACKNOWLEDGMENTS

This study was supported by JSPS KAKENHI Grant Number 24K08956.

## CONFLICT OF INTERST

The authors declare no conflicts of interest.

---

**Figure S1.**
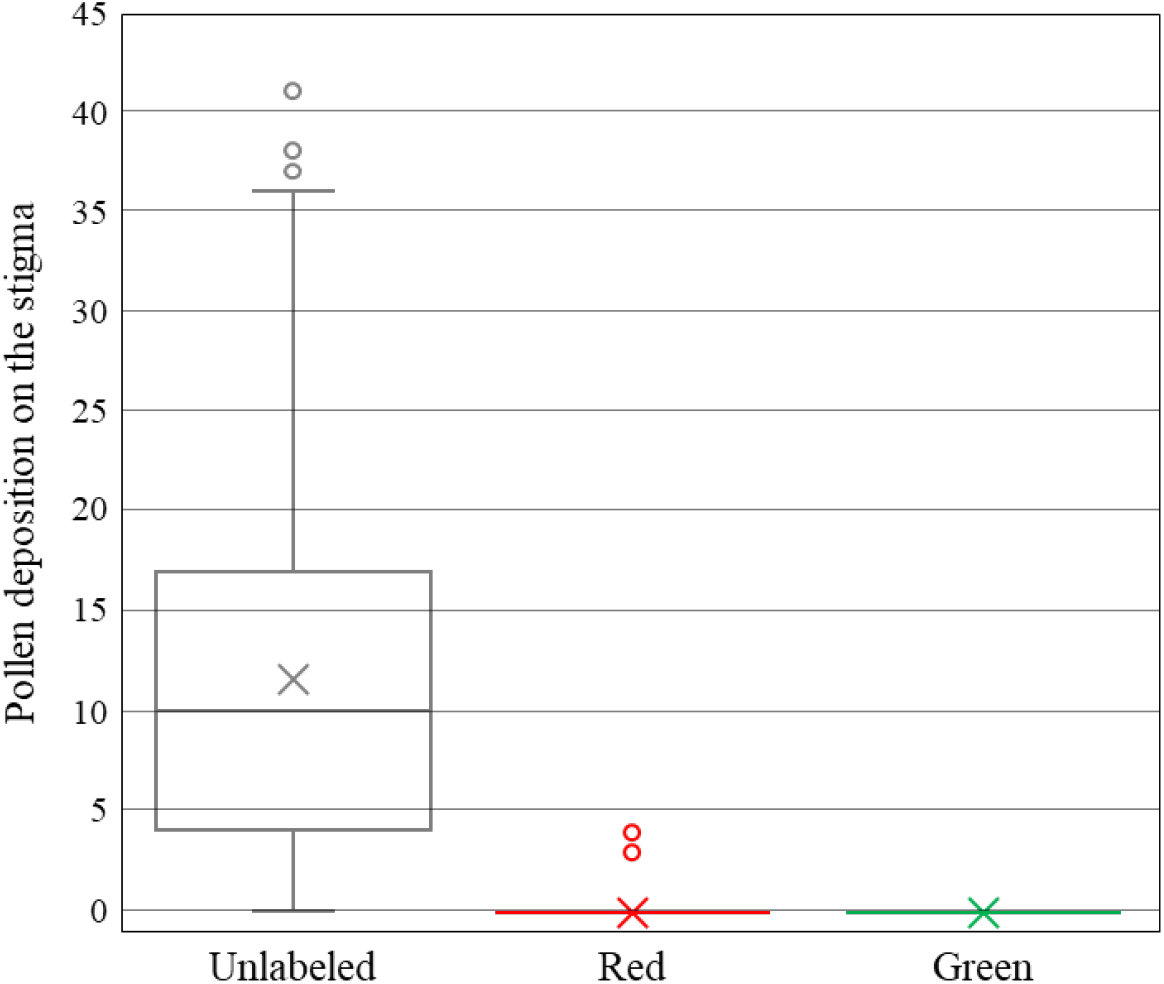
Pollen deposition on the stigmas of non-target flowers in field experiments. Unlabeled pollen represents a mixture of bud pollination and outcrossing with non-target flowers, whereas red- and green-labeled pollen represent outcrossing from the L- and M-anthers of the target flower, respectively. Boxplots show the median (horizontal line), interquantile range (box), whiskers extending to 1.5 × IQR, and outliers (points). Crosses indicate means.

## Graphical abstract

Heteranthery in *Commelina communis* integrates a temporally structured two-step selfing system: prior selfing via bud pollination followed by delayed selfing mediated by the medium-length stamen (M-anther). By functioning both as a pollinator reward and a mediator of delayed selfing, the M-anther links outcrossing with reproductive assurance within a single flower.

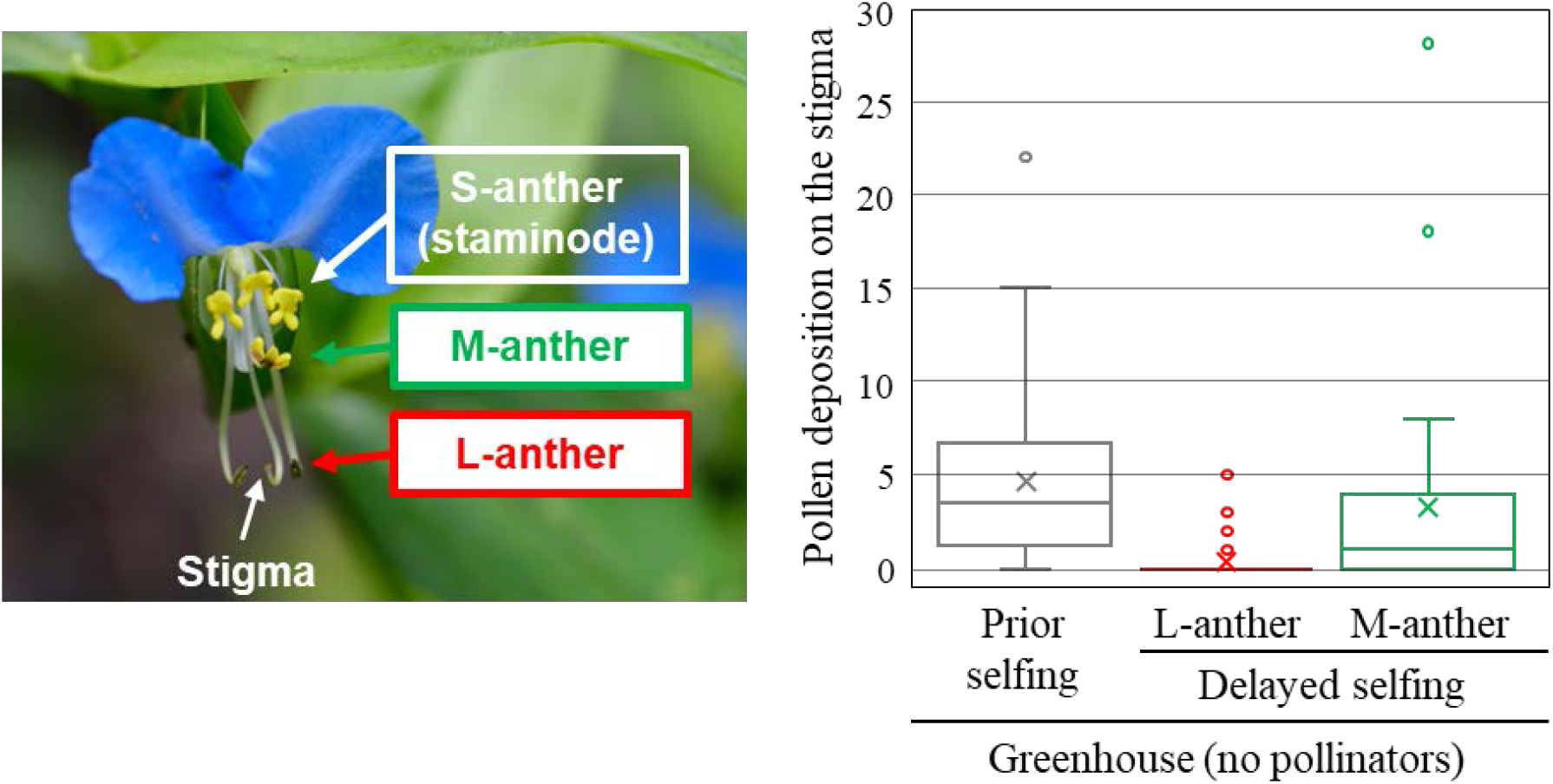

## Notes

### Competing Interest Statement

The authors have declared no competing interest.

